# Cross-species comparative hippocampal transcriptomics in Alzheimer’s disease

**DOI:** 10.1101/2021.06.09.447404

**Authors:** Marco Antônio De Bastiani, Bruna Bellaver, Giovanna Carello-Collar, Maria Zimmermann, Peter Kunach, Ricardo A. S. Lima-Filho, Stefania Forner, Alessandra Cadete Martini, Tharick A. Pascoal, Mychael V. Lourenco, Pedro Rosa-Neto, Eduardo R. Zimmer

## Abstract

Alzheimer’s disease (AD) is a multifactorial pathology, with most cases having a sporadic origin. Recently, knock-in (KI) models have been developed with the promise of resembling better sporadic human AD, such as the novel hAβ-KI mouse. Here, we compared hippocampal publicly available transcriptomic profiles of transgenic (5xFAD and APP/PS1) and KI (hAβ-KI) mouse models with early- (EOAD) and late- (LOAD) onset AD patients. Experimental validation of consistently dysregulated genes revealed four altered in mice (SLC11A1, S100A6, CD14, CD33, C1QB) and three in humans (S100A6, SLC11A1, KCNK). Additionally, the three mouse models presented more Gene Ontology biological processes terms and enriched signaling pathways in common with LOAD than with EOAD individuals. Finally, we identified 17 transcription factors potentially acting as master regulators of AD. Our cross-species analyses revealed that the three mouse models presented a remarkable similarity to LOAD, with the hAβ-KI being the more specific one.

## Introduction

Alzheimer’s disease (AD) is commonly categorized into early- (EOAD; <65 years old) and late-onset (LOAD; ≥65 years old) based on the age of clinical onset, the latter being the most prevalent form of the disease^1^. This broad definition includes the mendelian and non-mendelian EOAD, which confers an additional degree of heterogeneity to this group. Both forms of EOAD seem to present more prominent brain atrophy, glucose hypometabolism and increased tau PET uptake compared to LOAD^2–6^. However, a recent study demonstrated a similar biomarker profile between autosomal dominant EOAD and LOAD patients, supporting a shared pathobiological construct between both forms of the disease^7^.

Autosomal dominant inheritance mendelian EOAD accounts for less than 10% of the cases, but gene mutations found in these patients have been used for developing most AD models^1^. More specifically, mouse models overexpressing one or more mutations in the amyloid precursor protein (APP), presenilin 1 (PSEN1), and presenilin 2 (PSEN2) have dominated the AD experimental research. However, these models have undergone extensive scrutiny in the past years because these overexpression models present artifacts introduced by APP overexpression and do not resemble important aspects of LOAD^8,9,10^. Thus, a major challenge in the field has been the development of animal models that better recapitulate LOAD. The emergence of a new knock-in (KI) strategy for developing novel models expressing human APP with appropriate levels and cellular specificity seems to provide improved models for investigating sporadic AD^10–12^.

The investigation of core molecular programs shared by overexpression and KI models with human pathology may help determine to what extent animal models can resemble human disease. Additionally, the ability to recapitulate key molecular pathways activated or repressed in AD is crucial for model validity. Here, we aimed to ascertain the genes and pathways overlapping gene overexpression (5xFAD and APP/PS1) and KI (hAβ-KI) mouse models with EOAD and LOAD. With this in mind, we compared the hippocampal transcriptomic profiles of these animal models and AD and established the molecular similarity and specificity of mouse data to the human disease. We further employed a regulatory network-based approach to infer and investigate common master regulators between animal models and AD. Finally, we validated our transcriptomic exploratory findings in the hippocampus of APP/PS1 mice and *post-mortem* EOAD and LOAD individuals.

## Results

### hAβ-KI, 5xFAD, and APP/PS1 models exhibit more differentially expressed genes overlapping with LOAD than EOAD

We first investigated the genes that are differentially expressed between non-AD [wild type (WT, for animal models) and cognitively unimpaired individuals (for EOAD and LOAD)] and AD conditions. Differential expression analysis of the hippocampus of hAβ-KI, 5xFAD, and APP/PS1 mice and WT controls identified 1537, 3231, and 1768 differentially expressed genes (DEGs), respectively (unadjusted p-value < 0.05; **Figure 1B, C; Supplemental Figure S1, Supplemental Table 2**). Next, to investigate discrepancies in terms of gene expression between the three AD animal models and AD subtypes, we evaluated the overlap between DEGs in the different mouse models and human AD subtypes. We found that the hAβ-KI animals presented more DEGs overlapping with the 5xFAD mice than with APP/PS1 model [389 (25.3%) *versus* 235 (15.3%) DEGs, respectively; Chi-square adjusted p-value < 0.001; **Figure 1D**]. In addition, the comparison with AD human data demonstrated that hAβ-KI mice exhibited more DEGs in common with LOAD than with EOAD individuals (381 *versus* 164, respectively; **Figure 1E, F**). Despite being mouse models carrying familial AD-linked mutations, APP/PS1 and 5xFAD mice shared more DEGs with LOAD than with EOAD patients (**Figure 1E, F**). Moreover, considering the total of DEGs identified in the mouse models as reference (model-disease overlap), the intersection of DEGs between 5xFAD and EOAD was significantly higher (14%) than the overlap between both APP/PS1 (10.9%; Chi-square adjusted p-value = 0.0061) or hAβ-KI (10.7%; Chi-square adjusted p-value = 0.0054) with EOAD (**Figure 1E - right**). On the other hand, the overlap of LOAD DEGs with 5xFAD (28%) and with APP/PS1 (25.3%) mice was not significantly different from the overlap with hAβ-KI (24.8%; **Figure 1F - right**; Chi-square adjusted p-value = 0.123 and Chi-square adjusted p-value = 0.063, respectively). Interestingly, the hAβ-KI model shared 212 DEGs exclusively with LOAD, while only 98 with EOAD (**Figure 1B, C**). When we compared only the adjusted p-value DEGs [Benjamini & Hochberg (BH) > 0.1] we observed a similar profile/proportion of gene overlap, especially regarding the hAβ-KI model and LOAD (**Supplemental Figure S2**). Thus, we opted for using unadjusted p-values < 0.05 for subsequent functional enrichment analysis, which allows for exploratory cross-species analysis of core molecular programs that can be further validated. Only seven DEGs were found in common with all the mouse models and human AD. Among them, DEGs related to innate immune response (C1QB, CD33, CD14, and SLC11A1) were consistently upregulated across the groups, while those related to membrane potential regulation and neuropeptide production (KCNK1 and SST) were mostly downregulated. The S100A6 gene, found in neurons and some populations of astrocytes, was upregulated in all conditions (**Supplemental Table 2**).

**Figure 1.**
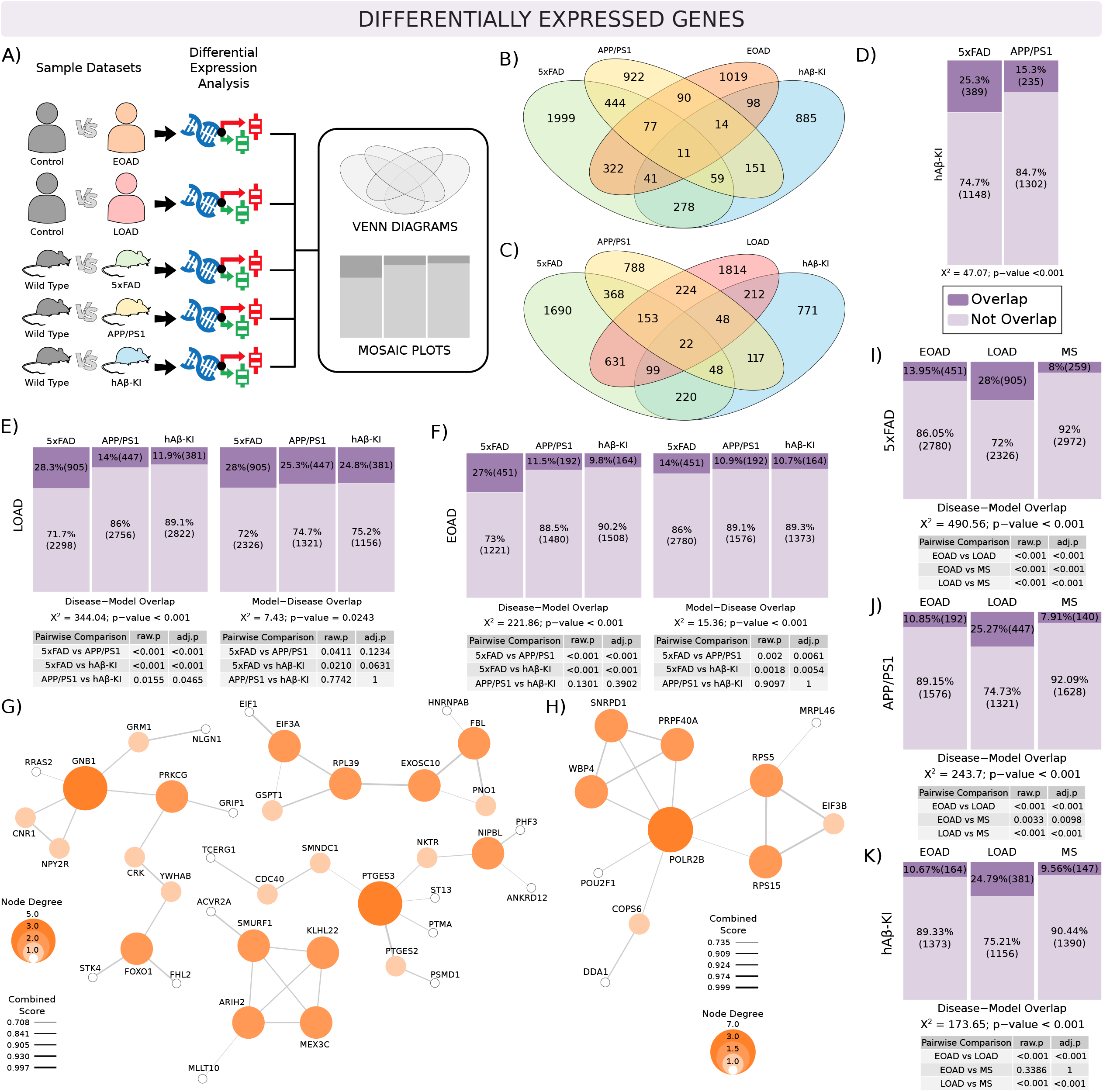
Shared DEGs among EOAD, LOAD and mouse models of AD. Schematic summary of samples used and analysis workflow to obtain DEGs **(A)**. Venn diagram showing DEGs overlap between EOAD **(B)** and LOAD **(C)** with 5xFAD, APP/PS1 and hAβ-KI mice. Mosaic plot of hAβ-KI, 5xFAD, and APP/PS1 overlap **(D)**. Mosaic plot of EOAD-model (left) and model-EOAD (right) DEGs overlap with 5xFAD, APP/PS1, and hAβ-KI mice **(E)**. Mosaic plot of LOAD-model (left) and model-LOAD (right) DEGs overlap with 5xFAD, APP/PS1 and hAβ-KI mice **(F)**. PPI network of DEGs exclusively shared between hAβ-KI mice and LOAD **(G)** or EOAD **(H)** patients. Mosaic plot of EOAD, LOAD or MS DEGs overlap with 5xFAD mice **(I)**. Mosaic plot of EOAD, LOAD or MS DEGs overlap with APP/PS1 mice **(J)**. Mosaic plot of EOAD, LOAD or MS DEGs overlap with hAβ-KI mice **(K)**. The size of red and yellow boxes reflects the proportion of overlapping and non-overlapping DEGs, respectively. Pearson’s Chi−squared test with Yates’ continuity correction was applied for the mosaic plot analysis. EOAD = early-onset Alzheimer’s disease; LOAD = late-onset Alzheimer’s disease; MS = multiple sclerosis; hAβ-KI = humanized amyloid-β knock-in. Genes with unadjusted p-value < 0.05 were considered as DEGs.

Because functional activity of proteins is highly dependent on their interactions with other proteins, understanding protein interactions is crucial to uncover their role. The protein-protein interaction (PPI) network of DEGs intersecting the hAβ-KI model and LOAD patients revealed that PTGES3, GNB1, ARIH2, SMURF1, EIF3A genes are hubs of the four clusters formed (**Figure 1G**). However, only two genes revealed in the PPI network – GNB1 and NKTR – were differentially expressed in the hAβ-KI model after BH adjustment. Considering the DEGs exclusively shared with the hAβ-KI model and EOAD patients, only 11 remained in the PPI network, and only the PRPF40A gene remained significant in hAβ-KI after multiple comparisons correction (**Figure 1H**). Next, we compared the DEGs of each mouse model with a database from multiple sclerosis (MS) patients to verify the specificity of these models for AD pathology. The three mouse models presented a greater overlap of DEGs with AD than with MS individuals (**Figure 1I, J; Supplemental Figure S3A, B**). Interestingly, no significant differences were observed in DEGs shared between hAβ-KI mice and EOAD (10.7%) or MS patients (9.6%; **Figure 1K; Supplemental Figure S3C**), while the overlap of hAβ-KI DEGs with LOAD was significantly higher than with both EOAD and MS (24.8%; Chi-square adjusted p-value < 0.001; **Figure 1K**; **Supplemental Figure S3C**).

To validate our exploratory transcriptomic findings, we performed qRT-PCR analyses of the seven DEGs overlapping between human AD and animal models. The validation was performed in APP/PS1, LOAD, and EOAD *post-mortem* hippocampus. Significant increases in of S100A6, C1QB, CD33, CD14, and SLC11A1 mRNA levels were observed in APP/PS1 mice compared to WT littermates (**Figure 2A**). Regarding the human hippocampal tissue validation, S100A6 was significantly increased in LOAD and EOAD compared to cognitively unimpaired individuals (**Figure 2B**). In addition, SLC11A1 expression increased, while KCNK significantly decreased in LOAD individuals. Although the other genes presented decrease/increase trends in AD patients compared to cognitively unimpaired individuals, they did not reach statistical significance.

**Figure 2.**
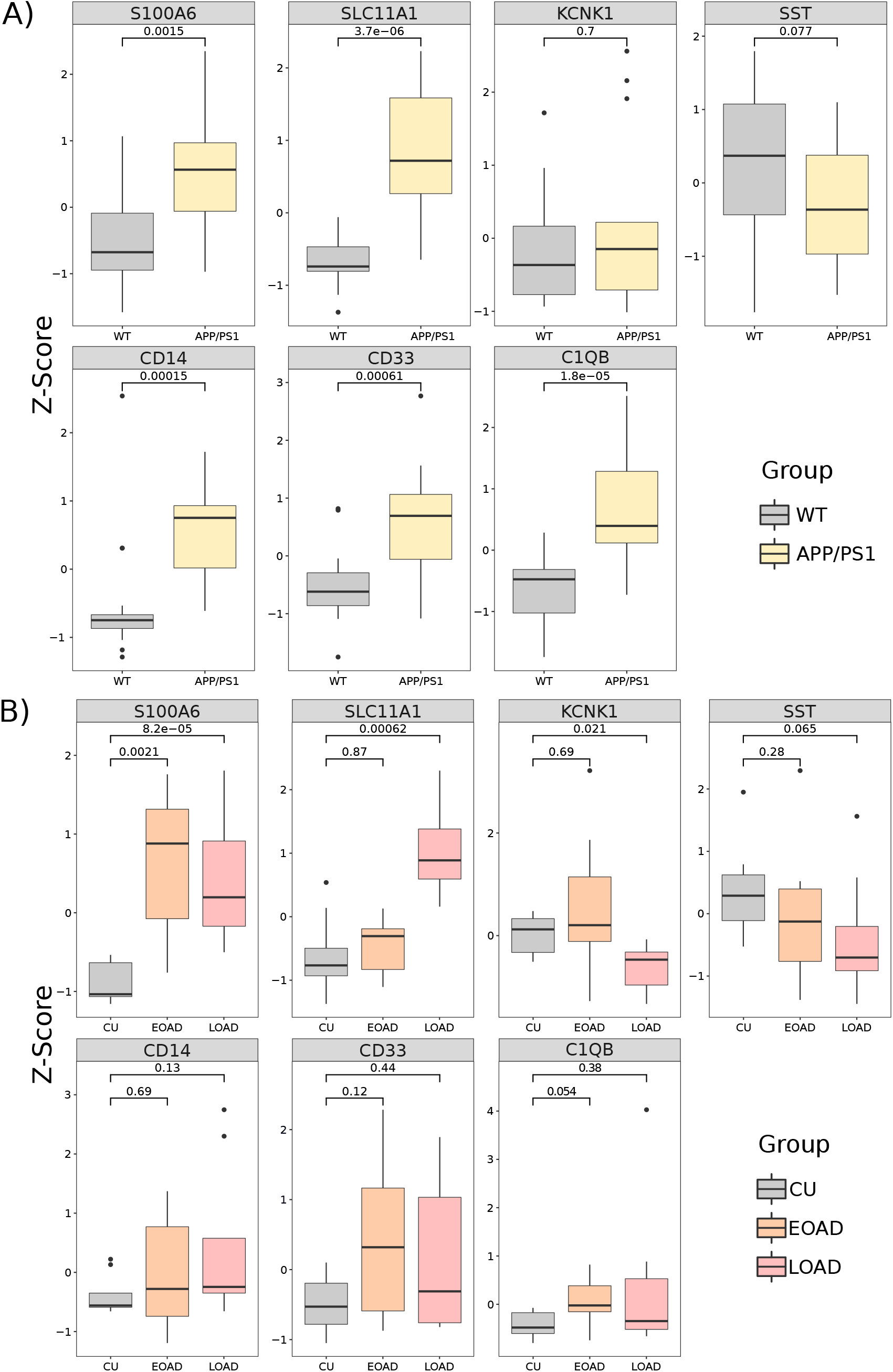
Experimental validation of exploratory transcriptomics findings. The expression seven genes overlapping between AD and animal models were evaluated by qRT-PCR in hippocampus from APP/PS1 (n = 14) and WT (n= 14) mice from two independent laboratories **(A)** and in cognitively unimpaired (CU, n = 9), early- (EOAD, n = 7) and late-onset Alzheimer’s disease (LOAD, n = 8) patients from the Douglas-Bell Canada Brain Bank **(B)**. Standard scores (z-score) of APP/PS1, EOAD and LOAD were compared for their difference from control using Wilcoxon.

### hAβ-KI mice and LOAD patients present higher similarities in functional changes

We next investigated the larger biological processes in which the DEGs found for each animal model/AD subtypes are involved. Functional enrichment analysis of gene ontology for biological processes (GOBPs) revealed that ∼ 92% of the GOBPs enriched in hAβ-KI mice overlap with enriched terms in LOAD patients (**Figure 3C, F - right**) in the model-disease approach, while the intersection with EOAD was markedly low (around 32%, **Figure 3B, E - right**). The remaining 8% of GOBP terms not shared between hAβ-KI mice and LOAD were related to RNA splicing and protein phosphorylation (**Supplemental Table 3**). Additionally, the hAβ-KI mice presented a higher number of enriched GOBP terms in joint with 5xFAD (79.2%) than with APP/PS1 (56.1%) model (**Figure 3D**; Chi-square adjusted p-value < 0.001). Surprisingly, 5xFAD and APP/PS1 shared more than 75% of their enriched GOBPs with LOAD patients (**Figure 3C, F - right**) and only ∼ 45% with EOAD (**Figure 3B, E - right**). However, we observed that about 50% of GOBPs overlap between EOAD, LOAD and 5xFAD, indicating a lack of disease subtype specificity for this model (**Supplemental Table 3**). The comparison among the GOBPs of the three models with MS demonstrated, however, greater specificity of these mouse models for AD (**Supplemental Figure S3**). **Figure 3G-I** shows that all mouse models evaluated presented less than 5% of enriched GOBP terms in common with MS.

**Figure 3.**
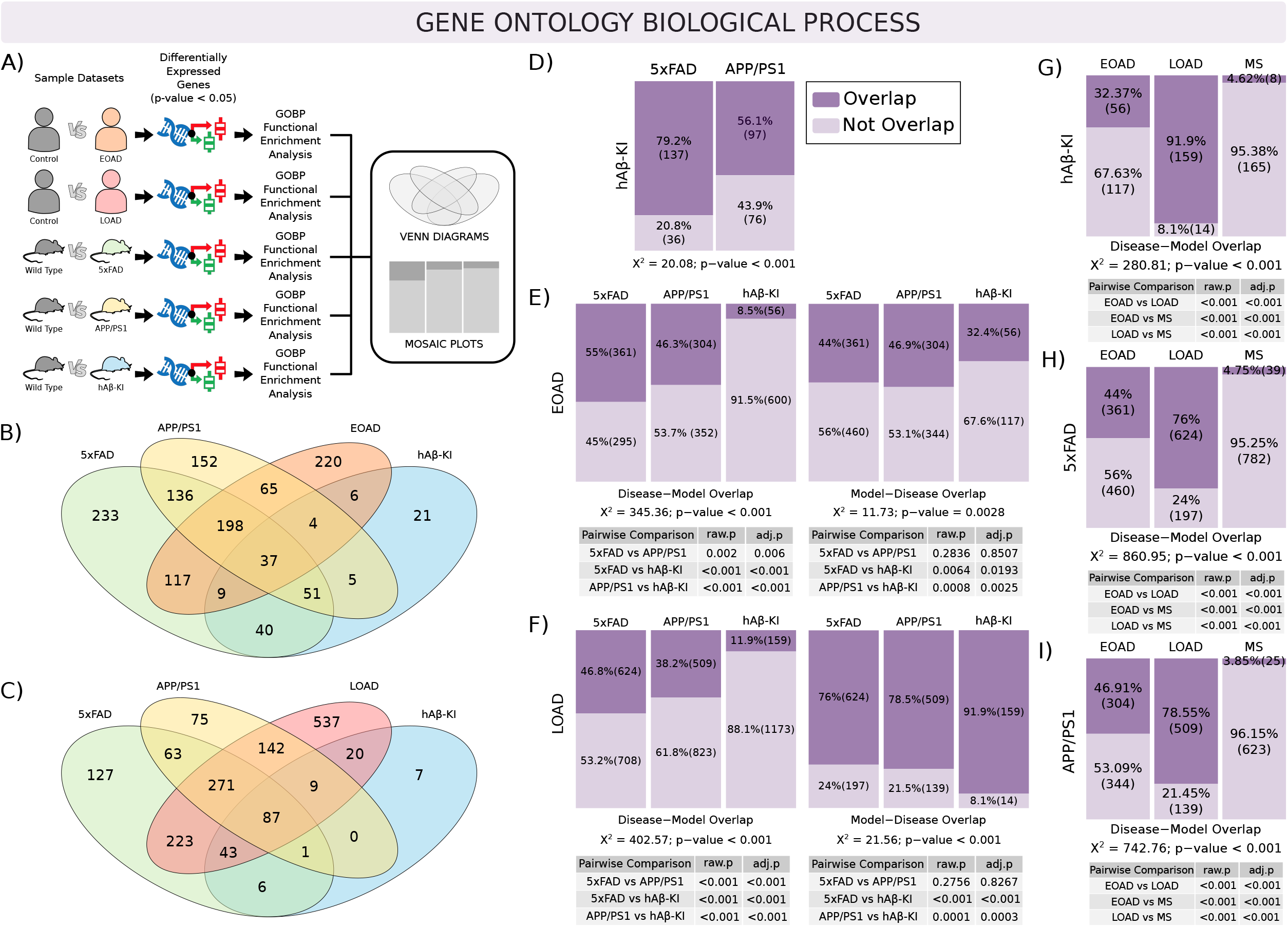
GOBP overlap among EOAD, LOAD, and mouse models of AD. Schematic summary of samples used and analysis workflow to obtain GOBPs **(A)**. Venn diagram of GOBPs intersections in EOAD **(B)** and LOAD **(C)** with 5xFAD, APP/PS1 and hAβ-KI mice. Mosaic plot of hAβ-KI-5xFAD and -APP/PS1 GOBP overlap **(D)**. Mosaic plot of EOAD-model (left) and model-EOAD (right) GOBP overlap with 5xFAD, APP/PS1, and hAβ-KI mice **(E)**. Mosaic plot of LOAD-model (left) and model-LOAD (right) GOBP overlap with 5xFAD, APP/PS1 and hAβ-KI mice **(F)**. Mosaic plot of EOAD, LOAD, or MS GOBP overlap with 5xFAD mice **(G)**. Mosaic plot of EOAD, LOAD, or MS GOBP overlap with APP/PS1 mice **(H)**. Mosaic plot of EOAD, LOAD, or MS GOBP overlap with hAβ-KI mice **(I)**. The size of the red and yellow boxes reflects the proportion of overlapping and non-overlapping GOBPs, respectively. Pearson’s Chi−squared test with Yates’ continuity correction was applied for the mosaic plot analysis. EOAD = early-onset Alzheimer’s disease; LOAD = late-onset Alzheimer’s disease; MS = multiple sclerosis; hAβ-KI = humanized amyloid-β knock-in.

### hAβ-KI mice present less GOBP terms intersection with EOAD patients compared to APP/PS1 and 5xFAD mice

As our initial analyses revealed several enriched GOBP terms, to better understand the global/general processes represent by them we computed the semantic similarity among GO terms. The union of enriched GOBP intersecting terms in the mouse models and human AD was generated to better visualize the common biological processes altered in each group. “Regulation of cytokine secretion” (13 nodes), “regulation of catabolic processes” (3 nodes), “NFκB signaling” (9 nodes), and “intracellular transport and secretion” (21 nodes) were among the GOBP terms enriched in the hippocampus of the three mouse models and EOAD patients (**Figure 4**; light gray circles). These terms are mainly related to cellular response to stressor agents, hormones and cytokines, immune response, and calcium homeostasis and transport. Interestingly, GOBP terms related to oxidative phosphorylation found in EOAD are only enriched in the APP/PS1 model (purple circles).

**Figure 4.**
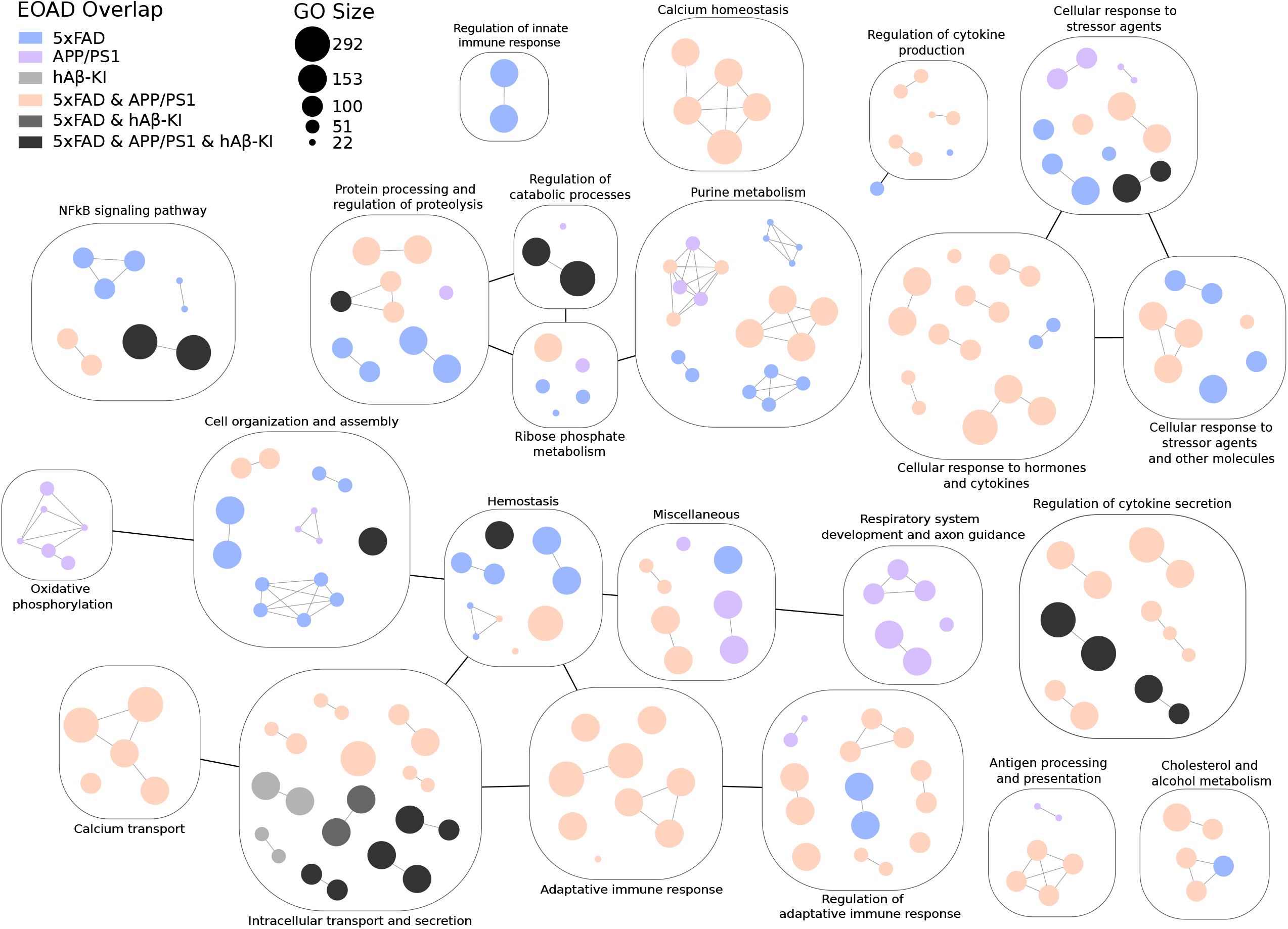
Nested networks of enriched GOBP intersections between EOAD and AD mouse models. Blue, purple and orange circles represent the overlap between EOAD and protein overexpression mouse models. Intersection among EOAD and hAβ-KI is represented by circles in shades of gray. Each box represents a cluster of GOBP terms grouped by semantic similarity and named manually according to its main biological role. Circle sizes represent the number of genes enriched in the GO term. EOAD = early-onset Alzheimer’s disease; MS = multiple sclerosis; hAβ-KI = humanized amyloid-β knock-in.

### hAβ-KI mice and LOAD patients exhibit greater overlap among the enriched GOBP terms

Similarly, we computed the semantic similarity among GOBP terms and compared the overlap with GOBP present in LOAD. **Figure 5** shows that hAβ-KI mice GOBP terms are more represented among the GOBPs in common with LOAD (**yellow, purple, and pink circles**) than with EOAD. “PI3K and NFkB signaling pathways” (20 nodes), “regulation of neuronal development” (19 nodes), “regulation of neurotransmitter and hormone secretion” (31 nodes), “regulation of immune response” (22 nodes), “antigen processing and presentation” (21 nodes) and “phospholipid and ribose phosphate metabolism” (10 nodes) are among the enriched biological processes shared between all three mouse models and LOAD (**pink circles**). Interestingly, “glucose, cholesterol and purine metabolism”, “regulation of MAPK cascade”, and “calcium homeostasis” only appeared enriched in 5xFAD and APP/PS1 mice (**gray circles**).

**Figure 5.**
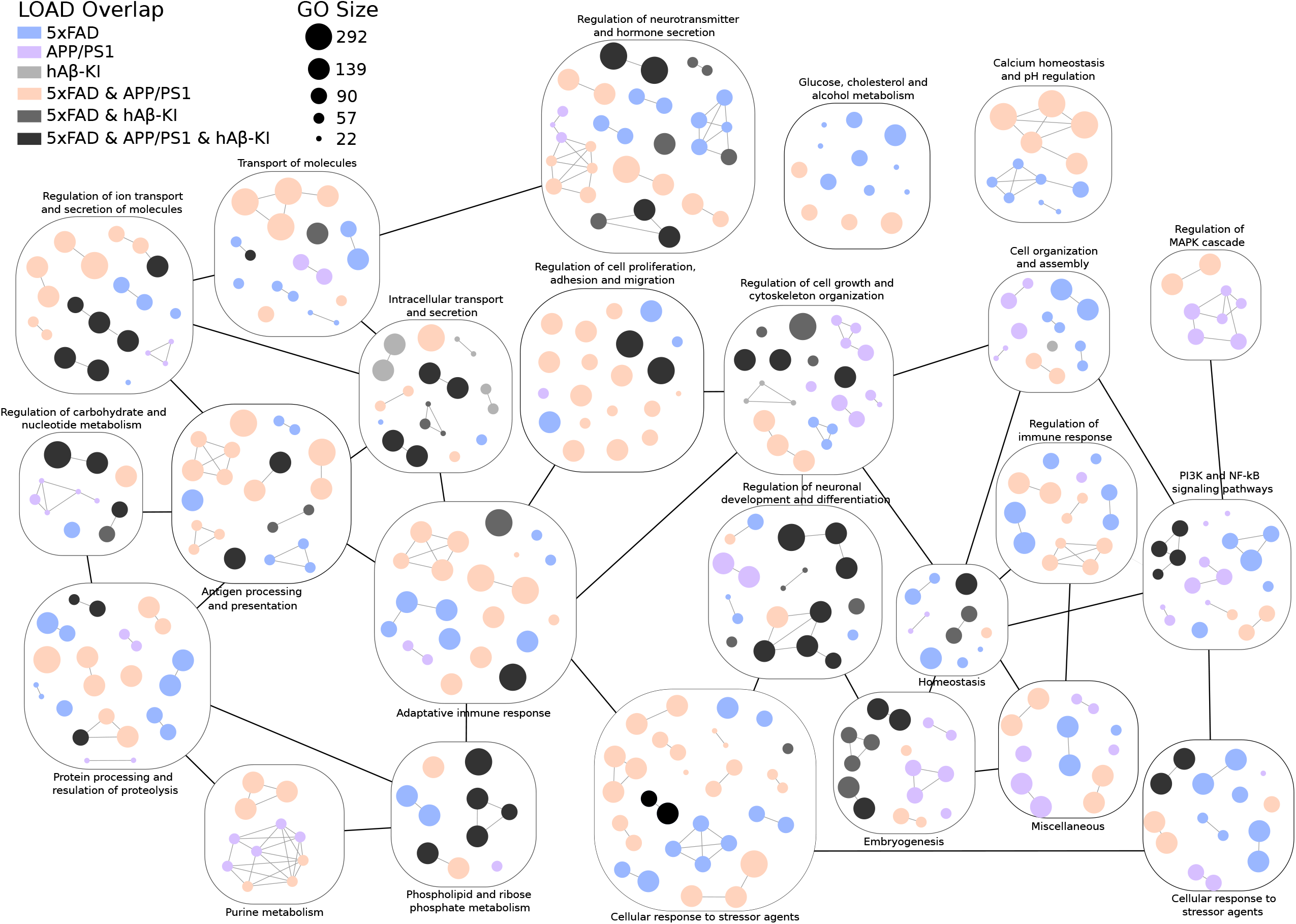
Nested networks of enriched GOBP intersections between LOAD and AD mouse models. Light purple, dark purple, and green circles represent all the GOBP terms overlap between LOAD and hAβ-KI. Intersection among LOAD and overexpression protein mouse models is represented by circles in shades of gray. Each box represents a cluster of GOBP terms grouped by semantic similarity and named manually according to its main biological role. Circle sizes represent the number of genes enriched in the GO term. LOAD = late-onset Alzheimer’s disease; MS = multiple sclerosis; hAβ-KI = humanized amyloid-β knock-in.

### Most of the hAβ-KI enriched KEGGs are shared with LOAD patients

The most affected pathways related to changes in transcriptome profile were identified using enrichment analysis of KEGG canonical pathways. **Figure 6A, B** shows that among the 10 KEGGs found significantly enriched in the hAβ-KI model, six were also identified in LOAD patients [“glutamatergic and GABAergic synapse” (adjusted p-value = 0.017), “calcium signaling” (adjusted p-value = 0.040), “Rap1 (adjusted p-value = 0.040) and Ras signaling (adjusted p-value = 0.040)” and “choline metabolism in cancer” (adjusted p-value = 0.041)], while only one was enriched in EOAD patients [“amyotrophic lateral sclerosis” (adjusted p-value = 0.040); **Supplemental Table 4**]. KEGG analysis also revealed that APP/PS1 was the mouse model that presented the highest percentage of pathway overlap with EOAD (36.7%; **Figure 6C**) and LOAD (73.3%; **Figure 6D**). Interestingly, most of the intersection between APP/PS1 and human AD is represented by KEGGs related to other neurodegenerative diseases (e.g. Parkinson’s disease, Huntington’s disease) or bacterial infection (e.g. *Escherichia coli, Salmonella* spp.; **Supplemental Table 4**). Despite PI3K pathway-related GOBPs appearing enriched in LOAD, they were only significantly altered in the hAβ-KI model (adjusted p-value = 0.040) in the KEGG analysis. “Rap1 and Ras signaling pathways” and “adhesion and apoptosis” were found consistently altered in the APP/PS1 and hAβ-KI models (**Figure 6H, I; Supplemental Table 4**). In addition, the 5xFAD was the mouse model that presented more enriched KEGG terms (**Figure 6G; Supplemental Table 4**). Similar to the GOBP enrichment analysis, 5xFAD presented more pathways shared with LOAD (56.9%) than with EOAD (13.8%; **Figure 6C, D**), which were mainly related to synaptic neurotransmission and insulin regulation (**Supplemental Table 4**). Additionally, alterations in apoptosis- and endocannabinoid system-related pathways were observed in the 5xFAD model and the human disease (**Supplemental Table 4**). Despite an overlap in KEGGs associated with inflammatory diseases and bacterial infections between 5xFAD and AD, well-studied signaling pathways in the context of inflammation were only observed in the 5xFAD model (*e*.*g*., TNF, Toll-like, NFκB).

**Figure 6.**
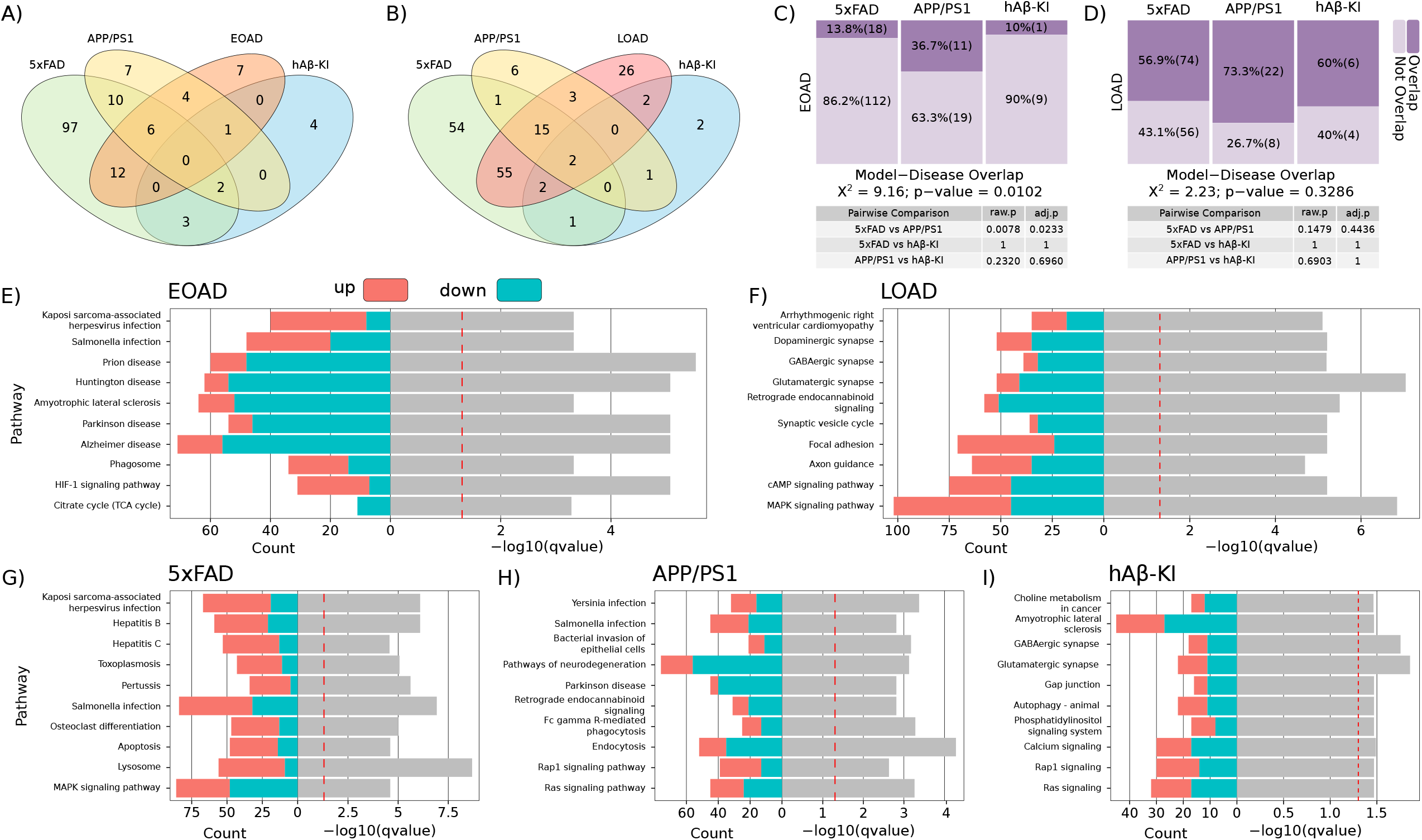
Functional enrichment analysis of KEGG terms in mouse models of AD and human disease. Venn diagrams showing the KEGGs overlap among EOAD **(A)** and LOAD **(B)** patients with 5xFAD, APP/PS1 and hAβ-KI mice. Mosaic plot of 5xFAD, APP/PS1 and hAβ-KI KEGGs overlap with EOAD **(C)** or LOAD **(D)**. Dual-plots showing the up- (red) and down- (blue) regulated KEGGs for EOAD **(E)**, LOAD **(F)**, 5xFAD **(G)**, APP/PS1 **(H)** and hAβ-KI **(I)**. The size of the red and yellow boxes reflects the proportion of overlapping and non-overlapping KEGGs, respectively. Pearson’s Chi−squared test with Yates’ continuity correction was applied for the mosaic plot analysis. The pathways in the dual-plots are represented in the Y axis; the left and the right part of the X axis corresponds to the number of genes enriched in the KEGG terms and their q-value (grey bars), respectively. EOAD = early-onset Alzheimer’s disease; LOAD = late-onset Alzheimer’s disease; MS = multiple sclerosis; hAβ-KI = humanized amyloid-β knock-in.

### EOAD and LOAD patients share two enriched master regulators with overexpressing and knock-in AD mouse models

To identify elements located in higher positions of the biological system hierarchy, we performed a master regulator analysis to determine transcription factors potentially driving the biological alterations observed in AD. In addition, we asked if these elements were also orchestrating the transcriptional profile changes in mouse models. Our analysis revealed a total of 95 master regulators enriched within DEGs emerging in at least one experimental group (**Figure 7A, B; Supplemental Table 5**). Interestingly, both 5xFAD and APP/PS1 mouse models presented more master regulators in common with LOAD than with EOAD, while hAβ-KI mice exhibited a similar overlap with both subtypes of human AD (**Figure 7A, B**). Among the 17 master regulators identified in ≥ 4 groups, only PARK2 and SOX9 were enriched in the three models and in the human disease (**Figure 7C**). Next, two-tail gene set enrichment analysis (GSEA) was performed to infer the activation state of each candidate. We observed that, while PARK2 is repressed, SOX9 is activated across the disease/animal model phenotypes evaluated (**Figure 7D**). In addition, FOXC2 and ZNF461 were identified exclusively in the hAβ-KI mice and LOAD individuals (**Supplemental Table 5)**. The three mouse models share eight enriched master regulators, most of which involved in regulating cell cycle and apoptosis (**Supplemental Table 5**). A second methodological approach to infer activation of transcription factors was also applied, and similar results were observed for the 17 master regulators candidates (**Supplemental Figure S4**). Finally, a comparison with a previously published study that investigated master regulators associated with AD showed that 11 out of 17 MR identified here were also enriched in that dataset (**Figure 7E**).

**Figure 7.**
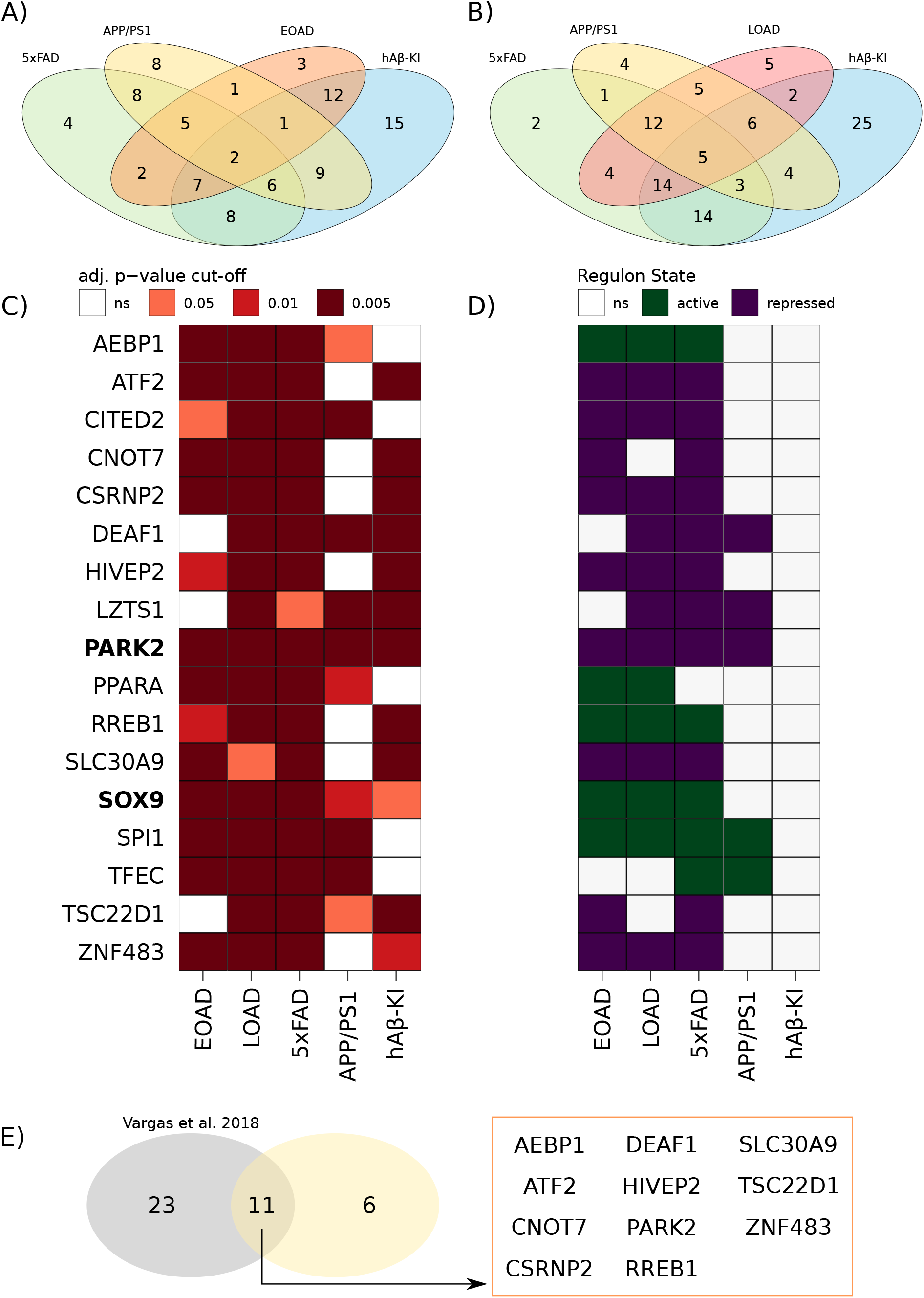
Master regulator analysis of mouse models of AD and human disease. Venn diagrams of EOAD **(A)** or LOAD **(B)** with 5xFAD, APP/PS1, and hAβ-KI MRs overlap. Tile-plot of regulons significantly enriched with DEGs in EOAD, LOAD, 5xFAD, APP/PS1 and hAβ-KI (only regulatory units significant in ≥ 4 contexts are represented) **(C)**. Tile-plot showing the activation state of the transcription factors acting as MR in EOAD, LOAD, 5xFAD, APP/PS1, and hAβ-KI **(D)**. Venn diagram showing the overlap between previously MR candidates found by Vargas et al. and the current study **(E)**. EOAD = early-onset Alzheimer’s disease; LOAD = late-onset Alzheimer’s disease; hAβ-KI = humanized amyloid-β knock-in; ns = non-significant.

## Discussion

In the past years, significant efforts of collaborative initiatives generated multiple mouse models with the promise of better recapitulating the phenotypic spectrum of sporadic LOAD. The evaluation of these animal models’ phenotype is a work in progress. Here, we evaluated hippocampal similarities and differences of three mouse models (APP/PS1, 5xFAD, and the novel hAβ-KI) at the transcriptional level. An exploratory cross-species comparative transcriptomics was also conducted to evaluate shared molecular core programs between these models and EOAD/LOAD cases. All mouse models showed more similarities to LOAD than to EOAD patients. The hAβ-KI mouse model presented not only a remarkable transcriptomic similarity for LOAD but also a specificity, and, surprisingly, the gene overexpressing models also better resemble LOAD than EOAD.

The specificity of the hAβ-KI model for LOAD was first observed in the exploratory DEG analysis, as hAβ-KI mice presented twice more DEGs exclusively overlapping with LOAD than with EOAD patients. Further PPI network analysis of these DEGs identified gene products that interact with each other to accomplish different biological functions. For example, we found clusters involved in ribosomal RNA processing, inflammatory response, and E3-ubiquitin ligase-related immune response, all phenomena well described in AD^13–15^. In line with this, Baglietto-Vargas and colleagues recently demonstrated that hAβ-KI mice presented a decreased production of the anti-inflammatory cytokines IL-2 and IL-10 compared to age-matched WT animals^11^. Importantly, a cluster of genes encoding G protein and G protein-coupled receptors (GPCRs) was also evidenced in our study. G proteins act as modulators or transducers in various transmembrane signaling systems, and GPCRs are implicated in multiple stages of AD pathogenesis^16^. Specifically, the glycogen synthase kinase 3-β (GSK3-β) is known for mediating tau phosphorylation and, consequently, being an active player in the development of tau pathology in AD^17^. Interestingly, this process seems to depend on the PI3K signaling activation^18^, a pathway significantly altered in the hAβ-KI mice. The clusters identified with our network analysis might shed light on the understanding of pathogenic mechanisms of LOAD that can be recapitulated by the novel hAβ-KI mouse model, facilitating the search for therapeutic targets.

Interestingly, only seven (C1QB, CD33, CD14, SLC11A1, KCNK1 and SST) out of 7868 DEGs identified in our transcriptomics analysis were shared among the mouse models, EOAD and LOAD patients. Multiple studies have already implicated these genes in AD pathophysiology. For instance, a decrease in somatostatin (SST) gene expression in AD brains has been previously reported^21^. Interestingly, SST was shown to be the most selectively enriched binder to soluble oligomeric Aβ in the human brain, influencing Aβ aggregation and masking the ability of widely-used antibodies to detect Aβ^22^. Nilsson and colleagues further demonstrated that modulation of the SST receptors Sst1 and Sst4 regulates neprilysin, the major Aβ-degrading enzyme^23^. The consistent findings observed in our study suggest that SST role in AD pathology seems to be conserved cross-species. Additionally, we found that C1QB, CD33, CD14, and SLC11A1 genes, all associated with immune responses, were upregulated in all analyzed mouse models, as well as in EOAD and LOAD patients. Accordingly, Gjoneska and colleagues previously pointed a conserved immunological basis for AD by comparing the CK-p25 mouse model with hippocampal human *post-mortem* tissue^14^. Our qRT-PCR analysis in APP/PS1 mice confirmed alterations in these genes. The analysis in the human brain only reached statistical significance for three of the seven genes – S100A6, SLC11A1, KCNK1 – but a trend was observed for SST and C1QB. This validation step indicates that our exploratory analysis provides meaningful biological information regarding AD pathology in the hippocampus.

The identification of transcriptomic changes at the pathway level has the potential to offer insights into the biological processes disturbed in AD. We observed that the hAβ-KI hippocampus presented an almost complete overlap of enriched hippocampal GOBP terms with LOAD, while only about one-third was shared with EOAD. This specificity for LOAD seems to be a unique and important feature of this novel KI model, as the 5xFAD and APP/PS1 mice also presented a significant overlap with EOAD. Despite that, the scanty overlap of GOBP terms among AD mouse models and MS confirmed that these models present transcriptomic features specific to AD rather than general alterations shared among other neurodegenerative diseases. In addition, this resemblance with AD appears to be a specific feature of rodent models carrying mutated human genes related to AD-associated amyloid pathology, such as APP and PSEN1, or humanized Aβ-KI. Indeed, Burns and colleagues observed that Tg4510 mice, which express a tau mutation found in familial frontotemporal dementia, presented the highest enrichment of genes in common with human ALS and Huntington’s disease rather than with AD^24^.

Impaired calcium (Ca^2+^) handling by neurons precedes the formation of amyloid plaques and neurofibrillary tangles and has been implicated in major molecular alterations underlying AD^25^. The association between PS1 mutations and altered Ca^2+^ signaling in neurons has been known for many years^26^. Early imbalance between excitatory/inhibitory (E/I) neurotransmission, with loss of neuronal network stability, is a well-attested phenomenon in AD^27,28^. The average incidence of seizure in AD patients is around 15%, which is seven-fold higher when compared to individuals without dementia^29,30^. Aβ is able to impair the long-term potentiation, promote depression of synaptic activity and alter the brain network^28^, and affect the E/I balance by impairing the inhibitory activity of the parvalbumin-expressing and SST interneurons^31,32^. Interestingly, modulation of interneuron function might improve brain rhythms and cognitive functions in AD^31,33^. Our study identified “calcium signaling” and “glutamatergic and GABAergic signaling” to be exclusively altered in the hAβ-KI mice and LOAD, pointing to this KI mouse model as an important tool to better understand calcium signaling and the E/I imbalance in AD.

Transcription factors play a key role in orchestrating phenotypic determination by regulating transcriptional targets that coordinate complex cellular processes^34,35^. We identified two transcription factors exclusively altered in the hAβ-KI model and LOAD individuals’ hippocampi: ZNF461 and FOXC2. ZNF461 roles in brain function have still been poorly explored; however, this transcription factor was identified among genes that represent a polygenetic risk for psychiatric disorders, and its alteration might contribute to cortical atrophy and changes in functional connectivity^36^. On the other hand, FOXC2 function in the brain is implicated in cell proliferation and invasion in glioblastoma^37^, in angiogenic processes during fetal brain development^38^, and is directly regulated by cyclin-dependent kinase 5 (Cdk5) phosphorylation to control peripheral lymphatic vase development^39^. Interestingly, it has been demonstrated that the deregulation of Cdk5 contributes to AD pathology preceding tau hyperphosphorylation and loss of synaptic proteins^40^. Several studies using organotypic hippocampal slices and primary neural cells exposed to Aβ showed increased p25 generation independently of APP overexpression^41–43^, suggesting that experimental validation of FOXC2 and Cdk5 in hAβ-KI mice is needed to understand the potential link among FOXC2, Cdk5, and AD pathology.

In the master regulator analysis, we identified SOX9 and PARK2 as transcription factors consistently activated in the mouse models and human AD. SOX9 is a key factor in the nervous system development, especially for astrocyte and oligodendrocyte cell fate specification^44–46^. Recently, Sun and colleagues identified SOX9 as an astrocyte-specific nuclear marker in the adult human and mouse brains, presenting a remarkable expression in the murine hippocampus and cortex compared to the cerebellum^47^. However, there are no studies to date linking this transcription factor with AD, and our results highlight a promising new target for investigation in AD. PARK2 encodes an E3 ubiquitin ligase, and its involvement in autosomal recessive parkinsonism is well established. Although less explored, its role in AD has already been demonstrated by computational^35,48^ and experimental^49–51^ approaches. Specifically, mitophagy failure, promoted by repression in PARK2 ability to stimulate PS1, was reported in cellular and animal models^49,51^. On the other hand, the overexpression of PARK2 promoted diminished brain accumulation of ubiquitinylated proteins, improved its targeting to mitochondria, and potentiated autophagic vesicle synthesis^50^. Our findings thus highlight the value of hAβ-KI, 5xFAD, and APP/PS1 mouse models to better understand these particularly underexplored aspects in AD.

Decades of use of animal models in AD research underline that each of them can mimic a slightly different aspect of the disease. Therefore, one could argue that animal models of AD should be selected according to the biological aspect aimed for investigation rather than be seen as a generic model. The clusterization by semantic similarity of enriched GOBP terms performed here allowed highlighted alterations specific to 5xFAD, APP/PS1 or hAβ-KI mouse models. For example, several GOBP terms associated with oxidative phosphorylation, purine metabolism, and the MAPK pathway were altered in 5xFAD and APP/PS1 mice but not in the hAβ-KI model. On the other hand, inflammatory-related processes seem to be a feature of AD pathology present in all three mouse models. Thus, the comparison of altered biological processes among mouse models and human AD presented here sheds light on the translational power of each animal model and helps to improve our mechanistic understanding of AD pathology.

### Limitations of the study

This study attempts to unveil core molecular functions in three AD animal models and compare them with human pathology transcriptomic profiles. To increase the sensibility for detecting biological processes and pathways, genes with unadjusted p-value < 0.05 were considered as DEGs. We experimentally validated the consistently altered genes found in our study in the APP/PS1 mouse model and human brain samples. However, the functional changes inferred from DEGs in our study should be further investigated in future studies.

## Methods

### Mouse Models Data Acquisition

RNA sequencing (RNA-seq) data from 5xFAD [4, 8 and 12 months-old, n = 23 hemizygous; 26 wild type (WT)] and hAβ-KI (22 months-old, n = 7 homozygous; 8 WT) AD mouse models (**Supplemental Table 1**) were obtained from AMP-AD Knowledge Portal (https://adknowledgeportal.synapse.org/) using *synapser* (version 0.7.64) and *synapserutils* (version 0.1.6) packages. Specifically, gene expression information was collected from https://www.synapse.org/#!Synapse:syn16798173 and https://www.synapse.org/#!Synapse:syn18634479 for 5xFAD and hAβ-KI models, respectively. APP/PS1 (8 and 12 months-old, n = 8 APP/PS1; 8 WT) mouse model RNA-seq data (**Supplemental Table 1**) was combined from two Gene Expression Omnibus (GEO) (https://www.ncbi.nlm.nih.gov/geo/) datasets [GSE149661^52^ and GSE145907] and downloaded through NCBI Sequence Read Archive using SRAToolKit (https://github.com/ncbi/sra-tools). After quality control evaluation, the following samples were removed: sample “67-2” from hAβ-KI; samples “466”, “305”, “456”, “497” from 5xFAD. Known phenotypic features of the three mouse models evaluated in this study are depicted in **Figure 8A**.

**Figure 8.**
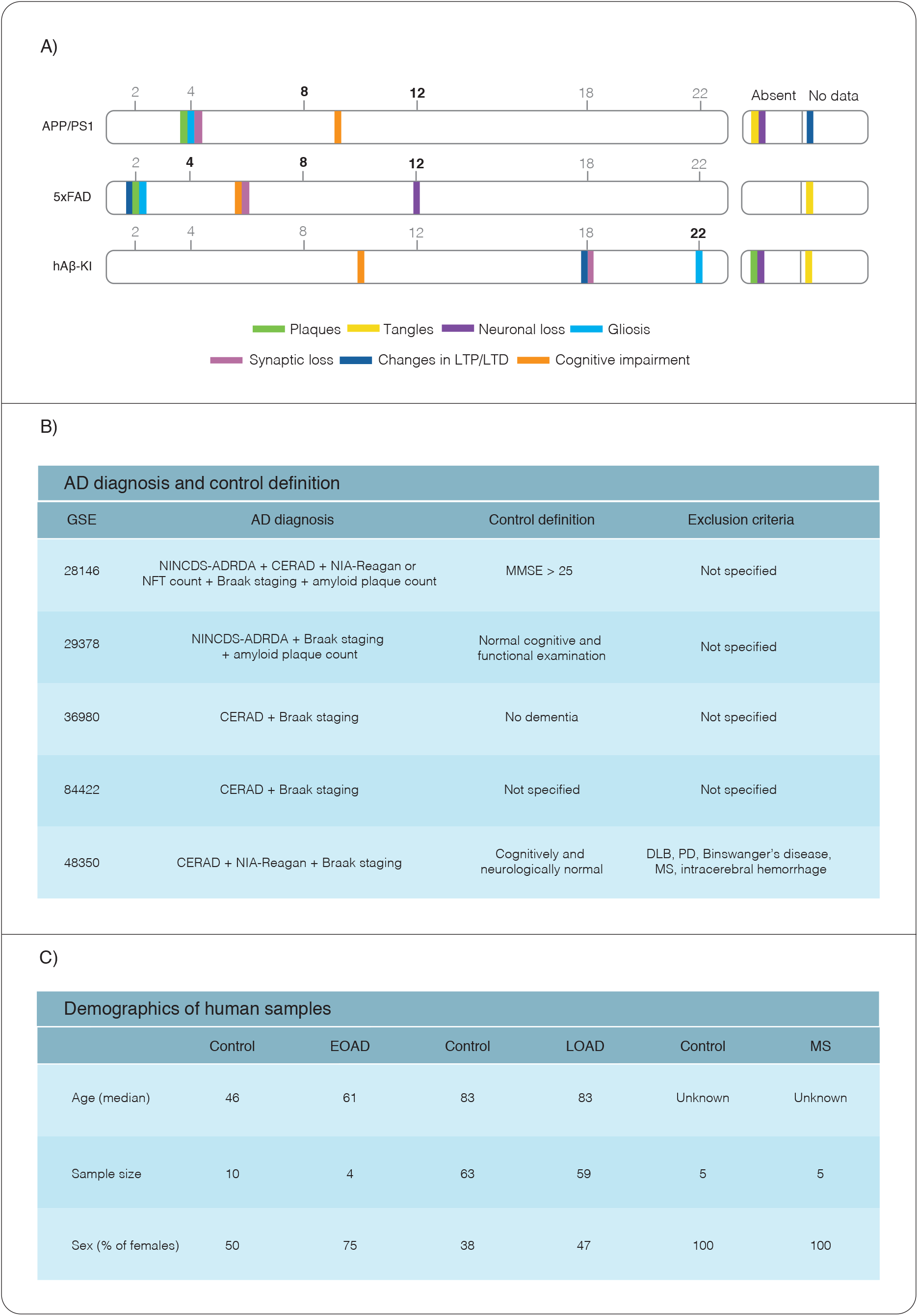
Phenotypic features of AD mouse models, AD diagnosis, control group definition, and demographics of human samples. APP/PS1 (top), 5xFAD (middle) and hAβ-KI mice (bottom) timelines showing specific phenotypes of each mouse model across different ages (in months (**A**). Numbers in bold correspond to the time points evaluated in this study. Data for APP/PS1 and 5xFAD mice were obtained at AlzForum. hAβ-KI information was taken from Baglietto-Vargas et al (Baglietto-Vargas *et al*., 2021). The individuals were diagnosed with AD according to established clinical criteria and/or neuropathological *post-mortem* evaluation in each selected GSE study **(B)**. Age, sample size and sex information are specified for EOAD, LOAD and MS patients and their age-matched controls **(C)**. NINCDS-ADRDA 1984 (McKhann *et al*, 1984); CERAD (Mirra *et al*, 1991); NIA-Reagan Institute criteria (Hyman, 1997); Braak staging (Braak & Braak, 1991); NFT = neurofibrillary tangles; MMSE = Mini-Mental State Examination; DLB = Dementia with Lewy bodies; PD = Parkinson’s disease; MS = Multiple Sclerosis. EOAD = early-onset Alzheimer’s disease; LOAD = late-onset Alzheimer’s disease.

### Human Data Acquisition

Human AD hippocampal processed microarray data of five studies were obtained from GEO repository [GSE28146^53^, GSE29378^54^, GSE36980^55^, GSE48350^56^ and GSE84422^57^], downloaded using GEOquery package (v2.56.0)^58^ and combined under common gene symbol annotations. Afterwards, batch correction was implemented using the sva package (v3.36.0) ^59^ and data was split into EOAD (age at death < 65, n = 4 EOAD; 10 cognitively unimpaired controls) and LOAD (age at death ≥ 65, n = 59 LOAD; 63 cognitively unimpaired controls) for further analyses (**Supplemental Table 1**). AD diagnosis, control definition and exclusion criteria of each GSE study are depicted in **Figure 8B**. Multiple sclerosis (n = 5 MS; 5 control) hippocampal RNA-seq data (**Supplemental Table 1**) was also obtained from GEO under the identifier GSE123496^60^ and downloaded through NCBI Sequence Read Archive. Sample demographics from EOAD, LOAD and MS individuals can be found in **Figure 8C**.

### RNAseq Processing

Raw data for each RNAseq dataset was downloaded and transcript alignment was performed using Salmon (v1.3.0) (Patro *et al*, 2017). Transcripts were mapped to genome using indexes derived from *Mus musculus* GRCm38 Ensembl build (ftp://ftp.ensembl.org/pub/release-96/fasta/mus_musculus) and *Homo sapiens* GRCH38 Ensembl build (ftp://ftp.ensembl.org/pub/release-96/fasta/homo_sapiens) for the mouse models and human data, respectively. Aligned reads were summarized using tximport (v1.12.3)^61^ and genes with minimum mean of counts per million cut-off < 2 were filtered out.

### Differential Expression Analyses

Differential expression was computed on processed microarray data using the limma package^62^ *lmFit* function to fit multiple linear models by generalized least squares. In addition, *eBayes* function was used to compute moderated t-statistics, moderated F-statistic and log-odds of differential expression by empirical Bayes moderation of the standard errors towards a common value. For RNA-seq datasets, processed expression data from each study was submitted to DESeq2 (v1.28.1)^63^ method using previously created tximport Summarized Experiment. Differential expression analysis (DEA) based on the Negative Binomial distribution was computed with the *DESeq* function followed by log fold change shrinkage with the *lfcShrink* function (shrinkage estimator type = “ashr”). Genes with unadjusted p-value < 0.05 were considered as DEGs. For information about BH adjusted p-values, see **Supplemental Table 2**.

Venn diagrams for DEGs were constructed using VennDiagram package (v1.6.20)^64^. The proportion differences between overlapped genes between models and diseases were computed by Pearson’s Chi−squared test with Yates’ continuity correction for count data, followed by post-hoc pairwise Bonferroni adjustment. We compared the overlap of DEGs considering (i) the model-disease, where we obtained the number/proportion of molecular alterations observed in the models that are associated with the disease and (ii) the disease-model where we explored how much of the total transcriptomic alteration of the disease each model captures. Both measures together can give a measurement of fitness for modeling the pathology.

### Protein-Protein Interaction Network Reconstruction

We used the Search Tool for the Retrieval of Interacting Genes/Proteins (STRING) Consortium to build protein-protein interaction (PPI) networks. STRING is a biological database and web resource of known and predicted PPI which contains information from numerous sources, including experimental data, computational prediction methods and public text collections. The construction of the PPI networks was implemented in R using the STRINGdb (v2.0.2), RedeR (v1.36.0) and igraph (v1.2.6) packages^65–67^. For the final networks, we retained only the edges with a combined interaction score > 0.7 from all sources and highly connected nodes for the final networks.

### Functional Enrichment Analyses

DEGs (unadjusted p-value < 0.05) from human or mouse model studies were submitted to Gene Ontology (GO) and Kyoto Encyclopedia of Genes and Genomes (KEGG) enrichment analysis using the clusterProfiler package (v3.16.1) *enrichKEGG* and *enrichGO* functions. The GO terms were clustered by semantic similarity using the *mgoSim* function from GOSemSim (v2.14.2) package^68^ (arguments measure = “Wang” and combine = NULL). The resulting similarity matrices were represented as GO networks using the RedeR (v1.36.0) package^65^ for interactive visualization and manipulation of nested networks. Clusters of GO terms obtained from GOSemSim algorithm were manually named for their biological interpretation. Venn diagrams for enriched GO/KEGG terms were constructed using VennDiagram (v1.6.20) package. Nested networks were constructed by maintaining only the intersecting GOBP terms among the mouse models and each human pathology (either EOAD or LOAD). Finally, Jaccard coefficient > 0.7 was used to filter out edges with low gene intersection between terms. The proportion differences between overlapped terms between models and diseases were computed by Pearson’s Chi−squared test with Yates’ continuity correction for count data, followed by post-hoc pairwise Bonferroni adjustment.

### Reverse Engineering of Transcriptional Network

The transcriptional network (TN) centered on transcription factors (TF) and their predicted target genes were inferred using a large cohort of neurologically and neuropathologically normal individuals (n = 122) obtained from GEO under the identifier GSE60862^69^. Herein, the terms “regulatory unit” or “regulon” are used to describe the groups of inferred genes and their associated TFs. RTN (v2.12.1) package was used to reconstruct and analyze TNs based on the mutual information (MI) using the Algorithm for the Reconstruction of Accurate Cellular Networks (ARACNe) method^70–72^. In summary, the regulatory structure of the network is derived by mapping significant associations between known TFs and all potential targets. To create a consensus bootstrap network, the interactions below a minimum MI threshold are eliminated by a permutation step and unstable interactions are additionally removed by bootstrap. Finally, data processing inequality algorithm is applied with null tolerance to eliminate interactions that are likely to be mediated by a third TF. The reference hippocampus TN was built using the package’s default number of 1000 permutations and 100 bootstraps (p-value < 0.001).

### Master Regulators Inference and Two-Tailed Gene Set Enrichment Analysis

Master regulator analysis (MRA) was employed for the MR inference^34^. MRA computes the statistical overrepresentation of DEGs (p-value < 0.05) obtained from DEA in the regulatory units of the reference TN. The regulons were considered altered in the disease if they presented (1) statistical enrichment of DEGs, (2) regulon size > 50 and (3) ≥ 80% of the queried case-control studies. Two-tailed Gene Set Enrichment Analysis (GSEA) was also performed using the RTN package (version 2.4.6, p-value < 0.05 and 1000 permutations). Briefly, Pearson’s correlation was used to split the regulatory units into positively (A) and negatively (B) associated targets. Afterwards, the phenotype association of each subgroup was tested using the GSEA^73^ statistics, resulting in independent enrichment scores for each subgroup. Finally, we tested the differential enrichment (ESA – ESB) considering the following desirable criteria for clear association: (1) a maximum deviation from zero near opposite extremes and (2) a good separation of the two distributions. Thus, a high negative differential score implies that the regulon is repressed in the disorder phenotype, while a high positive one indicates that the regulon is induced.

### Virtual Inference of Protein Activity by Enriched Regulon Analysis

The virtual inference of protein activity by enriched regulon analysis (VIPER) is another regulatory network-based approach to infer protein activity from gene expression profiles. Similar to MRA, VIPER systematically analyze the expression of the regulatory units previously identified by ARACNe algorithm. However, VIPER uses a fully probabilistic, yet efficient enrichment analysis framework based on analytic rank-based enrichment analysis. The analysis was implemented using the viper (v1.22.0) package in R^74^.

### Mice

Male and female APPSwe/PS1ΔE9 mice on a C57BL/6J (#005864) background were originally obtained from The Jackson Laboratories and bred at our animal facility. WT littermates were used as controls. All genotypes were confirmed before use and reconfirmed after tissue extraction. Animals were housed in groups of up to five per cage with food and water *ad libitum*, under a 12 h light–dark cycle, with controlled room temperature. For qRT-PCR experiments, the whole hippocampus of 14 APP/PS1 (12-16 months-old) and 16 WT mice (12-15 months-old) were used.

### Human tissue

Globally sampled hippocampal tissue from LOAD (n = 8, mean ± SD age = 79.5 ± 5.6, 2F/6M), EOAD (n = 7, mean ± SD age = 51.3 ± 5.6, 6F/1M) and cognitively unimpaired individuals (n = 9, mean ± SD age = 74.6 ± 8.9, 4F/5M) were obtained from the Douglas-Bell Canada Brain Bank with the approval of the scientific journal of the Brain Bank and the research ethics boards of the Douglas Institute (approval number : IUSMD20-02).

### RNA extraction and qRT-PCR

Total RNA from hippocampus of LOAD, EOAD and cognitively unimpaired individuals as well as APP/PS1 and WT mice were isolated using TRIzol Reagent (Invitrogen Carlbad). The concentration and purity of the RNA were determined spectrophotometrically at a ratio of 260/280. Then, 1 µg of total RNA was reverse transcribed using Applied Biosystems™ High-Capacity cDNA Reverse Transcription Kit (Applied Biosystems, Foster City, CA), according to manufacturer’s instructions. Real-time quantitative polymerase chain reaction (qRT-PCR) was performed on an Applied Biosystems 7900HT system with SYBR green master mix (Applied Biosystems). Target mRNA levels were normalized using -β actin gene (Actb) as housekeeper and cycle threshold (Ct) values were used to calculate fold changes in gene expression relatively to cognitively unimpaired individuals or WT mice using the 2^-ΔΔCt^. For the human RNA extraction, Direct-zolTM RNA Microprep from Zymo Research was used. Gene expression analysis for human samples was performed at the Institute for Research in Immunology and Cancer (IRIC) Genomics Core Facility, Université de Montréal. qRT-PCR analysis in APP/PS1 and WT mice was performed independently at Universidade Federal do Rio Grande do Sul and at Universidade Federal do Rio de Janeiro. Primers used for qRT-PCR and result tables are listed in **Supplemental Table 6 and 7**. Standard scores (z-score) of APP/PS1, EOAD and LOAD were compared for their difference from control using Wilcoxon test in R statistical environment.

## DECLARATIONS

### Ethics approval and consent to participate

Not applicable.

### Consent for publication

Not applicable.

### Availability of data and materials

All datasets used in this manuscript are publicly available in the referenced repositories. Further information will be readily available upon request.

### Author Contributions

MAB selected the databases and performed transcriptomics analysis. MAB, BB and GC-C analyzed and interpreted the data and conceived figures and tables. BB, GC-C, RASL-F, MVL, MZ and PK collected experimental data. MAB and BB wrote the manuscript. ERZ conceived and supervised the project. GC-C, TAP, SF, ACM, RASL-F, MVL, PR-N and ERZ, edited and revised the manuscript for intellectual content.

## Acknowledgments

The results published here are partially based on data obtained from the AD Knowledge Portal (https://adknowledgeportal.synapse.org/). The IU/JAX and UCI MODEL-AD Center was established with funding from The National Institute on Aging (U54 AG054345-01 and AG054349).

## Supplemental Data Legends

**Supplemental Figure S1** | **DEGs in mouse models of AD and human disease**. Volcano plots and pie charts showing the up- and down-regulated genes in 5xFAD **(A)**, APP/PS1 **(B)**, hAβ-KI **(C)**, EOAD **(D)**, LOAD **(E)** and MS individuals **(F)**. Red points in volcano plots correspond to a DEG when compared to control. In pie charts, pink and blue colors correspond respectively to the up- and down-regulated DEGs. EOAD = early-onset Alzheimer’s disease; LOAD = late-onset Alzheimer’s disease; MS = multiple sclerosis; hAβ-KI = human amyloid-β knock-in. Genes with unadjusted p-value < 0.05 were considered as DEGs.

**Supplemental Figure S2** | **Intersection of DEGs between animal models and AD subtypes**. Venn diagram showing DEGs overlap between EOAD **(A)** and LOAD **(B)** with 5xFAD, APP/PS1 and hAβ-KI mice. Mosaic plot of EOAD-model **(A)** and model-EOAD **(B)** DEGs overlap with 5xFAD, APP/PS1 and hAβ-KI mice. Genes with *BH adjusted p-value < 0*.*1* were considered as DEGs in the mosaic plots.

**Supplemental Figure S3** | **Shared DEGs and enriched GOBP among mouse models of AD and human disease**. Venn diagrams showing the DEGs (left) and GOBP terms (right) overlap among 5xFAD **(A, D)**, APP/PS1 **(B, E)** and hAβ-KI **(C, F)** mice with EOAD, LOAD and MS individuals. EOAD = early-onset Alzheimer’s disease; LOAD = late-onset Alzheimer’s disease; MS = multiple sclerosis; hAβ-KI = human amyloid-β knock-in. Genes with unadjusted p-value < 0.05 were selected as DEGs.

**Supplemental Figure S4** | **VIPER analysis of EOAD, LOAD and AD mouse models**. Plot of inferred regulon activity showing the up- (reddish) and down- (bluish) regulated transcription factors in each group: EOAD **(A)**, LOAD **(B)**, 5xFAD **(C)**, APP/PS1 **(D)** and hAβ-KI **(E)**. EOAD = early-onset Alzheimer’s disease; LOAD = late-onset Alzheimer’s disease; hA β-KI = human amyloid-β knock-in. *p-value < 0.05

**Supplemental Table 1** | **Detailed description of transcriptomics datasets and samples**. Sample information for EOAD, LOAD, APP/PS1, 5xFAD, hAβ-KI and MS from repository datasets.

**Supplemental Table 2** | **Full tables of differentially expression analyses**. Differential expression results from EOAD, LOAD, APP/PS1, 5xFAD, hAβ-KI and MS.

**Supplemental Table 3** | **Full tables of Gene Ontology functional enrichment analyses and semantic similarity results**. GO biological processes functional enrichment analyses and semantic similarity results for EOAD, LOAD, APP/PS1, 5xFAD, hAβ-KI and MS.

**Supplemental Table 4** | **Full tables of functional enrichment analyses of KEGG terms**. Functional enrichment analyses results for EOAD, LOAD, APP/PS1, 5xFAD, hAβ-KI and MS.

**Supplemental Table 5** | **Table of results from master regulator analysis (MRA) and two-tailed gene set enrichment analysis (GSEA)**. MRA and two-tailed GSEA analyses results for EOAD, LOAD, APP/PS1, 5xFAD, hAβ-KI and MS.

**Supplemental Table 6** | **Table of oligonucleotide primers for qRT-PCR**. Primer sequences of selected mRNA for human and mouse experiments.

**Supplemental Table 7** | **Table of qRT-PCR results (z-scores)**. Standard scores (z-score) of APP/PS1, EOAD and LOAD computed from the 2^-ΔΔCt^.

**Supplemental Table 8** | **Tables of expression data**. Processed expression/count tables for each dataset used in the study.

**Supplemental Table 9** | **Tables of sample metadata**. Compiled tables of metadata information of each dataset used in the study.

**Supplemental Table 10** | **Tables of gene annotation**. Compiled tables of gene annotation for each dataset used in the study.

