## Supplemental Table 6 for "Cross-species comparative hippocampal transcriptomics in Alzheimer’s disease"

**Supplemental Table 6.** Oligonucleotide Primers for Real-Time RT-PCR

| mRNA target | Forward (5’-3’) | Reverse (5’-3’) |
| --- | --- | --- |
| Human |  |  |
| ACTB | TGACATTAAGGAGAAGCTGTGCTAC | ACTTCATGATGGAGTTGAAGGTAGT |
| C1QB | GGCTTCCAGGGCTGGCTGGAG | TCCCGATTCACCTTTGGGGCC |
| SST | CCCCAGACTCCGTCAGTTTCT | CATTCTCCGTCTGGTTGGGT |
| CD14 | CTGGAACAGGTGCCTAAAGGAC | GTCCAGTGTCAGGTTATCCACC |
| CD33 | TGTTCCACAGAACCCAACAA | GGCTGTAACACCAGCTCCTC |
| SLC11A1 | TCAAACTTCTCTGGGTGCTGCTCT | AGAGCAGATTGAATGCAATGGCCG |
| S100A6 | AAGCTGCAGGATGCTGAAAT | CCCTTGAGGGCTTCATTGTA |
| KCNK1 | GGGTGTGATGCTCCGTAGTC | ATATTGGGAGCCCCAGGCTT |
| Mouse |  |  |
| ACTB | CTTCTTTGCAGCTCCTTCGT | ATATCGTCATCCATGGCGAAC |
| C1QB | TCTGGGAATCCACTGCTGTC | AGACCTCACCCCACTGTGTC |
| SST | CTGCGACTAGACTGACCCAC | GAAACTGACGGAGTCTGGGG |
| CD14 | GAAGCAGATCTGGGGCAGTT | CGCAGGGCTCCGAATAGAAT |
| CD33 | GGTCAAAGTTCAGTGCCTTCCTC | TTGAGCATTTAAAACCCCAAAGC |
| SLC11A1 | TACCAGCAAACCAATGAGGA | CCTGGGGAAGATCTTAGCATAGT |
| S100A6 | CGCTTCTTCTAGCCCAGTCAT | ACTGGATTTCACCGAGAGAGG |
| KCNK1 | TCACATCATGGAGCACGACC | AGTCAAGGCTCTGCCTCTTG |
